# IAP retrotransposons contribute to the transcriptional diversity of the murine placenta

**DOI:** 10.64898/2026.01.27.702056

**Authors:** Samuele M. Amante, Maria L. Vignola, Cyril Pulver, Tessa M. Bertozzi, Anne C. Ferguson-Smith, Marika Charalambous, Miguel R. Branco

**Affiliations:** Blizard Institute, Faculty of Medicine and Dentistry, QMUL, London, UK; Centre for Epigenetics, QMUL, London, UK; Department of Medical and Molecular Genetics, Faculty of Life Sciences and Medicine, King’s College London, London, United Kingdom; School of Life Sciences, Swiss Federal Institute of Technology Lausanne (EPFL), CH-1015, Lausanne, Switzerland; Department of Genetics, University of Cambridge, Cambridge, UK; Whitehead Institute for Biomedical Research, Cambridge, MA, USA

## Abstract

Transposable elements (TEs) have made important contributions to the evolution of the placenta, and are argued to have played a role in the wide inter-species diversification of this critical developmental organ. Co-option of TEs by host genomes has led to the genesis of important placental genes, as well as trophoblast-specific gene regulatory elements. But whilst multiple TE subfamilies have been shown to act as transcriptional enhancers in early mouse trophoblast development, it remains unclear to what extent TEs regulate placental gene expression after the establishment of a functional fetal-maternal interface. Here, we characterised the TE regulatory and transcriptional landscape in mouse placenta and gauged their evolutionary dynamics through a comparative approach. We found that overall, the gene regulatory potential of TEs is greatly diminished in differentiated mouse trophoblast when compared to their stem cell counterpart. However, evolutionarily young intracisternal A particle (IAP) elements are highly expressed in the placenta and create several alternative, placenta-specific transcriptional start sites for protein-coding genes. IAP elements that are active in the placenta drive species-specific expression of associated genes and display wide genetic diversity between mouse strains. These putative co-option events are therefore evolutionarily recent and may represent a prime example of how TE activity can drive fast placental evolution.

## Introduction

The placenta stands out as one of the most diverse and fast-evolving organs within therian species^1^. Evidence suggests that immunological and resource allocation conflicts between mother and fetus led to a genetic and epigenetic arms race that predominantly acted by altering the physiology and morphology of the feto-maternal interface^2^. Transposable elements (TEs) – mobile genetic elements that can quickly generate novel coding and regulatory sequences within a genome – are among the key genetic drivers in placental evolution^3,4^. Most notably, TEs have been co-opted multiple times in different evolutionary lineages to form syncytin genes, which encode for retroviral envelope proteins that mediate trophoblast cell-to-cell fusion^5^. Other TE-derived genes with known roles in the placenta include *Peg10*^6^, *Peg11*^7^ and supressyn^8^.

TEs have also made a widespread contribution to pregnancy evolution as gene regulatory elements. The transcriptional promoters of TEs, such as the long terminal repeats (LTRs) of endogenous retroviruses (ERVs) and the 5’ UTR of LINE elements can create alternative transcriptional start sites for host genes, leading to the generation of TE-gene chimeras. Numerous examples have been uncovered in human placenta, including tissue-specific promoters for *CYP19A1*, *NOS3* and *PTN*^9^. The same non-coding TE sequences can also act as distal enhancers to fine tune gene expression. Their activity is determined by the opposing actions of activating transcription factors and repressive KRAB zinc finger proteins, as well as the underlying 3D chromatin structure^10^. We and others have identified a large array of putative TE-derived enhancers in mouse and human trophoblast, some of which were shown to play important roles in gene regulation and cellular phenotypes^11–15^. Finally, a large proportion of long non-coding RNAs have originated from TEs, including the HERVH-derived *UCA1* gene that promotes proliferation of human trophoblast stem cells (TSCs)^16^.

The house mouse (*Mus musculus*) is a popular model to study placental development and function, and has been pivotal for demonstrating the extent by which placental dysfunction impacts embryonic development^17^. The mouse is also an attractive model for studying TE co-option in mammals, as it provides the possibility to test the impact of TEs on organismal phenotypes. Additionally, the rodent lineage has seen recent waves of TE expansion, including several ERV subfamilies that remain mobile^18^, which creates an opportunity to study particularly recent or even ongoing co-option events. However, most knowledge of TE co-option in mouse trophoblast is restricted to the stem cell state. ERVs from the RLTR13D5 and RLTR13B-related subfamilies bear hallmarks of transcriptional enhancers in TSCs in vitro and in vivo, and can have dramatic effects on the expression of nearby genes^11,12^. Upon in vitro differentiation of TSCs, the regulatory activity of these ERV subfamilies appears to be dramatically reduced^19^, but it is not known whether other TEs become active as promoters and/or enhancers in differentiated trophoblast in vivo. We therefore know remarkably little about the potential of TEs to regulate gene expression during the vast majority of mouse placental development. Notably, the evolutionarily young IAP family of ERVs has been proposed to play regulatory roles in the mouse placenta^20^, but current evidence is scarce.

Here we investigated the contribution of TEs as regulatory elements in the mouse placenta in vivo. We find a scarcity of TE-derived enhancers in differentiated trophoblast, but uncover a role for IAPs in creating tissue- and species-specific alternative promoters. We posit that young IAPs generate genetic and epigenetic diversity that creates opportunities for co-option as gene regulatory elements within the mouse placenta.

## Results

### Differentiated mouse trophoblast harbours few TE-derived enhancers

To assess how the landscape of gene regulatory TEs changes upon differentiation of mouse trophoblast, we performed chromatin profiling (CUT&Tag and/or ATAC-seq) and RNA-seq on TSCs and E14.5 placenta (Figure 1A). We used a well-studied TSC line (GFP TSC)^21^, as well as a line we derived from the same strain of mice used for collecting placentas (B6 TSC), which expressed key TSC markers when grown in stem cell conditions, and makers of syncytiotrophoblast and trophoblast giant cells after 4 days of in vitro differentiation (Figure S1A). For in vivo tissue, we used whole placentas, as well as sorted placental cells using reporter transgenes to separate trophoblast (marked with tdTomato) from extraembryonic mesoderm (marked with GFP)^22^. Cell type deconvolution based on single-nuclei RNA-seq (snRNA-seq) data^23^ confirmed that the tdTomato fraction was overwhelmingly made up of trophoblast cells (92%), and suggested that the same was true of whole placenta samples (83% trophoblast; Figure S1B).

**Figure 1.**
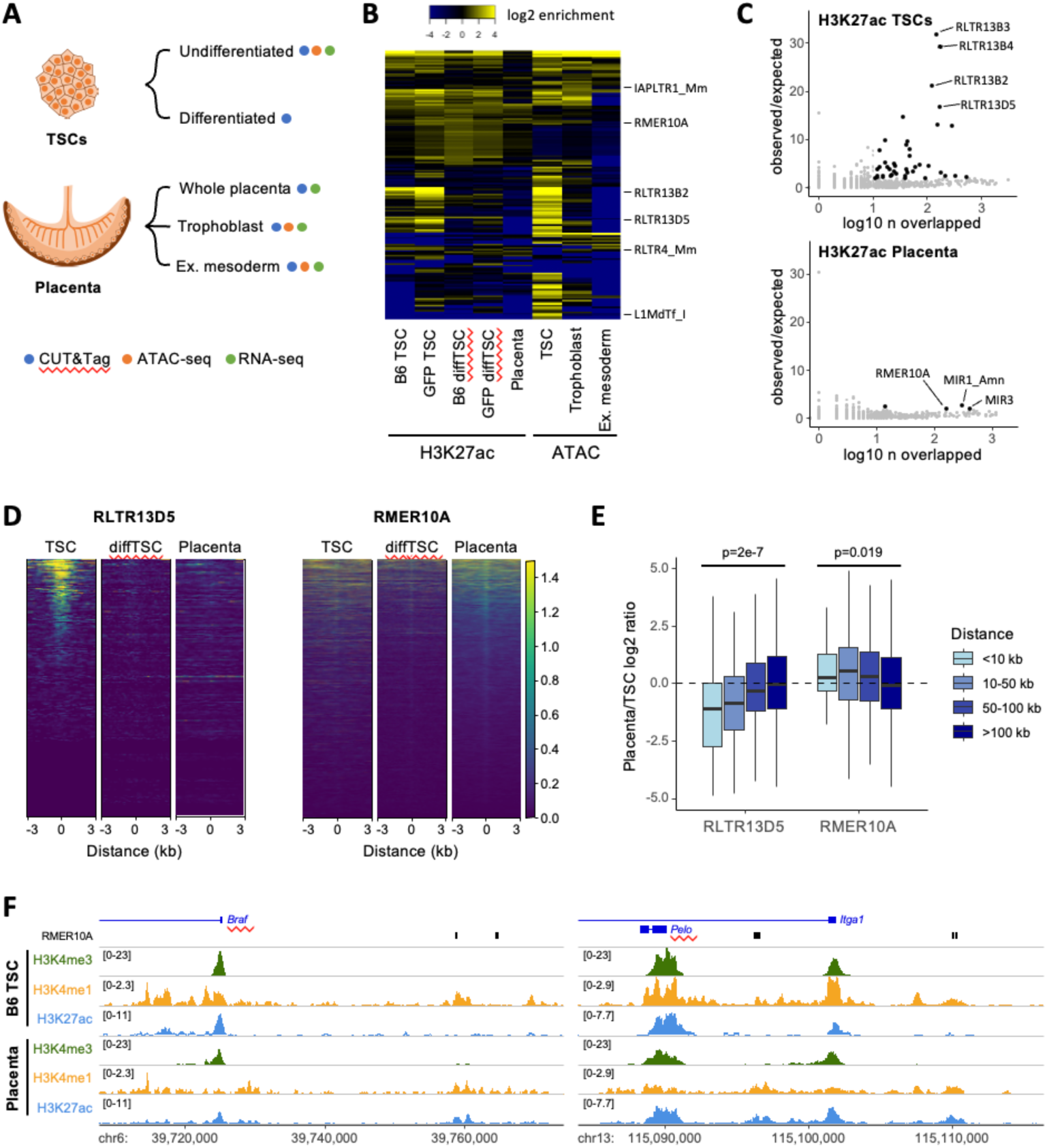
The mouse placenta has few TEs harbouring gene regulatory signatures. A) schematic of the profiling experiments performed/analysed. B) TE subfamilies displaying enrichment (observed/expected) for elements overlapping ATAC-seq or H3K27ac CUT&Tag peaks in at least one dataset (diffTSC: differentitated TSCs). C) TE subfamily H3K27ac enrichment in TSCs or placenta (black data points are classified as significantly enriched). D) H3K27ac CUT&Tag signal at RLTR13D5 or RMER10A elements (TSCs are B6). E) Relative expression (Placenta/TSCs) of genes containing TSSs within the indicated distances from an RLTR13D5 or RMER10A element. F) Examples for loci displaying placenta-specific enrichment of H3K27ac at RMER10A elements.

Using CUT&Tag and ATAC-seq data, we aimed to identify TE subfamilies that bear classic hallmarks of active promoters (open chromatin, H3K4me3, H3K27ac) and/or enhancers (open chromatin, H3K27ac, H3K4me1). We used a custom pipeline that selects TE subfamilies bearing more elements overlapping CUT&Tag/ATAC-seq peaks than expected by chance. This revealed highly contrasting profiles between undifferentiated and differentiated trophoblast, with a large number of TE subfamilies enriched for active/open chromatin in TSCs, whereas in vivo placenta/trophoblast had far fewer enriched subfamilies and smaller enrichment values (Figure 1B,C and S1C; Table S1). As expected from past work^11,12,19^, epigenetically active TE subfamilies in TSCs included RLTR13D5 and multiple RLTR13B-related subfamilies. In vitro differentiation of TSCs for four days was sufficient to diminish the enrichment for active chromatin marks at these subfamilies, as previously suggested^19^. We also inferred the cis-regulatory action of TE subfamilies in specific trophoblast cell types by analysing snRNA-seq data using the craTEs tool^24^. Whilst craTEs could accurately predict the *cis*-regulatory activities of RLTR13D5 and RLTR13B-related subfamilies in E6.5-E8.5 extraembryonic ectoderm (where the TSC niche sits)^25^, in the mature placenta only one TE subfamily (RLTR13D3A) was predicted to be active in a single trophoblast cell type (Figure S1D). Moreover, analysis of ENCODE H3K27ac ChIP-seq data (which closely correlates with our own data; Figure S1E,F) suggests that the mouse placenta has comparable or lower levels of TE regulatory activity to other somatic tissues (Figure S1G; Table S1). For example, testes have a far more striking enrichment for several TE subfamilies, including the well-characterised RLTR10B (Figure S1G,H)^26,27^. This contrasts with observations in human trophoblast^13–15,28^, where differentiated cells maintain a high level of putative regulatory activity at TEs (Figure S1G). Co-option of TEs as regulatory elements within the mature placenta may therefore not be as widespread across species as commonly thought.

We identified one TE subfamily (RMER10A) that displayed putative regulatory activity only in the placenta and not in TSCs, although the level of H3K27ac enrichment was far lower than what is seen for TEs that are active in TSCs (Figure 1D). Genes proximal to RMER10A elements enriched for H3K27ac displayed higher expression in placenta relative to TSCs (Figure 1E), supporting a potential role as distal enhancers. The expression differences were relatively mild when compared to the the effects that TSC-active TEs have on nearby genes (Figure 1E), suggesting a minor regulatory effect of RMER10A elements. Nonetheless, we could find clear examples of elements with an active enhancer-like profile (H3K4me1 + H3K27ac) close to genes relevant for placentation (Figure 1F). This includes *Itga1*, which is important for human trophoblast invasion^29^ and is thought to play similar roles in mouse^30^, as well as *Braf*, which is essential for mouse extraembryonic development^31^.

### IAP elements are highly expressed in differentiated trophoblast

We noted that several evolutionarily young TE subfamilies (IAPs, L1s, RLTR4) were enriched for open chromatin and/or H3K4me3 in trophoblast (Table S1). However, few uniquely mapped reads can be assigned to young TEs, limiting a full appreciation of epigenetic patterns therein. We therefore remapped our H3K4me3 data allowing for random allocation of non-unique reads and assessed the overall enrichment pattern over different young TE subfamilies. We found a strong enrichment for H3K4me3 at the 5’ ends of full-length elements from some IAP and L1 subfamilies, as well as RLTR4 elements (Figure 2A, S2A). Enrichment at IAPLTR1_Mm and L1Md_Tf/Gf subfamilies was specific to trophoblast, whereas RLTR4 elements were also enriched in extraembryonic mesoderm (Figure S2B). Similar patterns of enrichment are seen using uniquely mapped reads, although they are less pronounced (Figure S2C).

**Figure 2.**
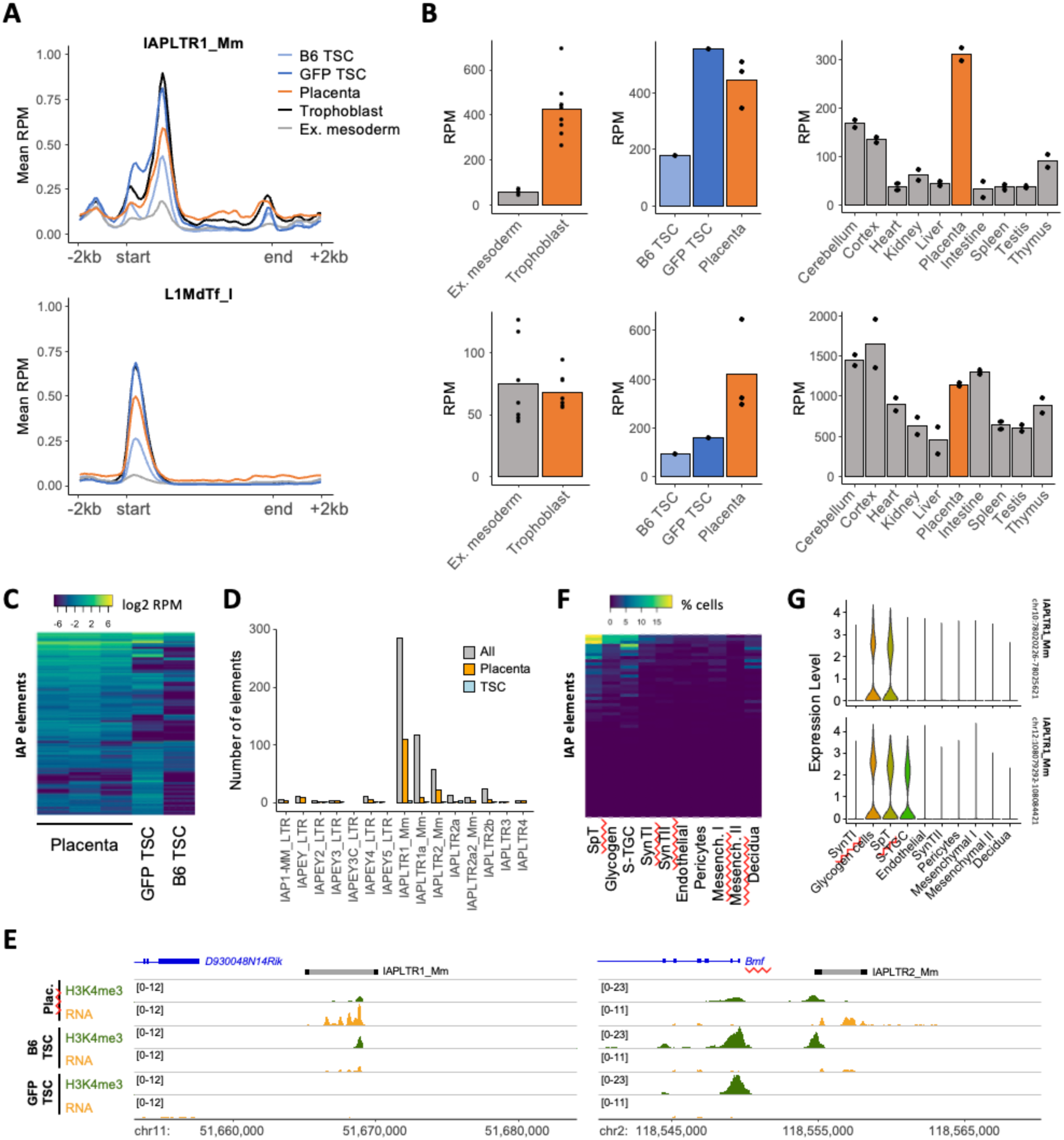
IAP elements are highly expressed in the placenta. A) Mean H3K4me3 **CUT<u>&</u>Tag** signal over IAPLTR1_Mm or L1MdTf_l elements, allowing for random assignment of non-unique reads. B) Expression of IAPLTR1_Mm (top) or L1MdTf_l (bottom) subfamilies in our own or ENCODE datasets, with non-unique reads assigned using **TEtranscrjpts. C)** Expression of individual IAP elements (unique reads only) in placenta or TSCs. D) Number of elements from each IAP subfamily expressed in all samples, or exclusively in placenta or TSCs. F) Percentage of cells expressing a given IAP element (unique reads) in clusters from snRNA-seq placenta data. G) Expression of two IAP element examples in placenta snRNA-seq data. E) Examples of two IAP loci showing placenta-specific expression.

To assess whether H3K4me3 enrichment was associated with TE expression, we first quantified subfamily-wide expression using TEtranscripts^32^ and SQuIRE^33^. Congruent with the H3K4me3 patterns, IAPLTR1_Mm expression was high in trophoblast (both undifferentiated and differentiated) but not extraembryonic mesoderm (Figure 2B, S2D). Indeed, analysis of ENCODE data showed that the placenta stands out as a tissue with high expression of IAPLTR1_Mm elements (Figure 2B, S2D). In contrast, expression of L1MdTf_I elements was not as pronounced in the placenta when compared to other tissues (Figure 2B, S2D). We then analysed the expression of individual loci using uniquely mapped reads. ERVs were defined using a consolidated annotation merging LTRs and internal fragments, and intragenic TEs were excluded to avoid confounding transcription originating from genes. This analysis revealed that expression over full-length elements was only robustly and unambiguously detected for IAP elements, but not L1 or RLTR4 elements (Figure S2E).

Focusing on IAPs, we found that there is a large number of elements that are specifically expressed in the placenta, especially when compared to low-passage TSCs of the same genetic background (B6 TSC; Figure 2C-E, Table S2). Expressed IAP elements were predominantly of the IAPLTR1_Mm subfamily (Figure 2D) and arranged in a proviral configuration (Figure S3A). A large proportion contained a known deletion characteristic of the IΔ1 subtype (Figure S3A,B), which is overrepresented in mutational IAP insertions^18^. Analysis of placental snRNA-seq data^23^ further showed that IAP expression is mainly restricted to spongiotrophoblast, glycogen cells and sinusoidal trophoblast giant cells (Figure 2F,G). Expression in spongiotrophoblast and glycogen cells is consistent with reports showing IAP expression at the junctional zone of the placenta, where these cell types reside^34,35^. To identify candidate transcription factors that might drive cell-specific expression, we performed binding motif analysis at IAP LTRs. As recently reported, we found binding motifs for CDX2 and TBX20 that may drive IAP expression in early trophoblast development^36^. In spongiotrophoblast, the most promising candidate was MITF, which is highly expressed in this cell type^23^ and whose binding motif is present in the most active IAP subfamilies in the placenta (Figure S3C).

We then asked whether IAP expression was associated with tissue- and locus-specific DNA methylation differences. Using published whole-genome bisulphite sequencing data^37^, we found that DNA methylation levels at IAPs were overall lower in the placenta when compared to brain, with H3K4me3-marked IAPs standing out as being particularly hypomethylated relative to other elements (Figure S3D). Notably, IAP elements have been shown to display large variation in DNA methylation across individuals from the same litter^38,39^. We found that 32% of our H3K4me3-marked IAPLTR1_Mm elements (compared to 2% of unmarked elements) were previously classified as variably methylated across littermates in B and T cells^38^. We therefore asked whether they remained variably methylated in the placenta. Bisulphite analyses showed that, despite displaying lower DNA methylation relative to tail biopsies, most IAPs analysed remain variably methylated in the placenta (Figure S3E). These data suggest that IAP expression may be variable across placentas, and thus any putative downstream effects may lead to inter-individual phenotypic differences.

### IAP elements generate placenta-specific gene isoforms

Given the enrichment for H3K4me3 at IAPs and young L1s in the placenta, we hypothesized that they might impact the expression of nearby genes. This could be by providing transcriptional start sites (TSSs) that generate chimeric TE-gene transcripts, or through more distal regulatory action, for which both mouse and human L1s have the potential^40–43^. To assess whether genes proximal to H3K4me3-marked elements are associated with increased gene expression, we first compiled lists of IAPs and L1s marked by H3K4me3 in both placenta and TSCs (‘common’) or only in placenta. We then quantified gene expression differences between placenta and TSCs, and found that genes close to IAPs or L1s that are active only in the placenta were preferentially expressed in this tissue (Figure 3A, S4A). Genes within 10 kb of these elements are also more highly expressed in the placenta relative to other somatic tissues, and specifically within the trophoblast fraction of the placenta (Figure 3B, S4B). In contrast, genes close to IAPs and L1s that are active in both placenta and TSCs displayed no expression differences (Figure 3A, S4A).

To test whether IAPs and L1s generate alternative TSSs in the placenta, we performed transcriptome assembly using our RNA-seq data, then selected for transcripts that initiate at IAPs or L1s and that incorporate exons from known genes. Using these criteria, we found that placental RNA samples harboured 52-66 IAP-associated transcripts, whereas TSC samples had only 11-23, which is consistent with the increase in IAP expression in the placenta. In contrast, there was no difference in the number of L1-associated transcripts between placenta (71-79) and TSCs (51-82). After stringent manual curation of merged transcript isoforms in placenta, we identified 27 high-confidence chimeric transcripts involving IAP elements and only 3 involving L1s (Table S3). The majority of these transcripts were only detected in placenta and not TSCs (Table S3, Figure 3C). Amongst them were two non-canonical isoforms of *Suv39h2* (Figure 3C), which encodes for a key H3K9 methyltransferase that controls trophoblast stem cell fate in rat^44^. *Suv39h2* transcriptional initiation switches dramatically from the canonical promoter in TSCs to IAP-based promoters (9-14 kb upstream) in the placenta (Figure 3C,D), but with no evidence that this leads to the production of a larger protein (Figure S5). Other IAP-associated genes with reported roles in trophoblast function were *Arhgef18*, a guanine exchange factor that is required for mouse syncytiotrophoblast development^45^, and *Acly*, which regulates histone acetylation in human trophoblast and may affect placental development^46^.

**Figure 3.**
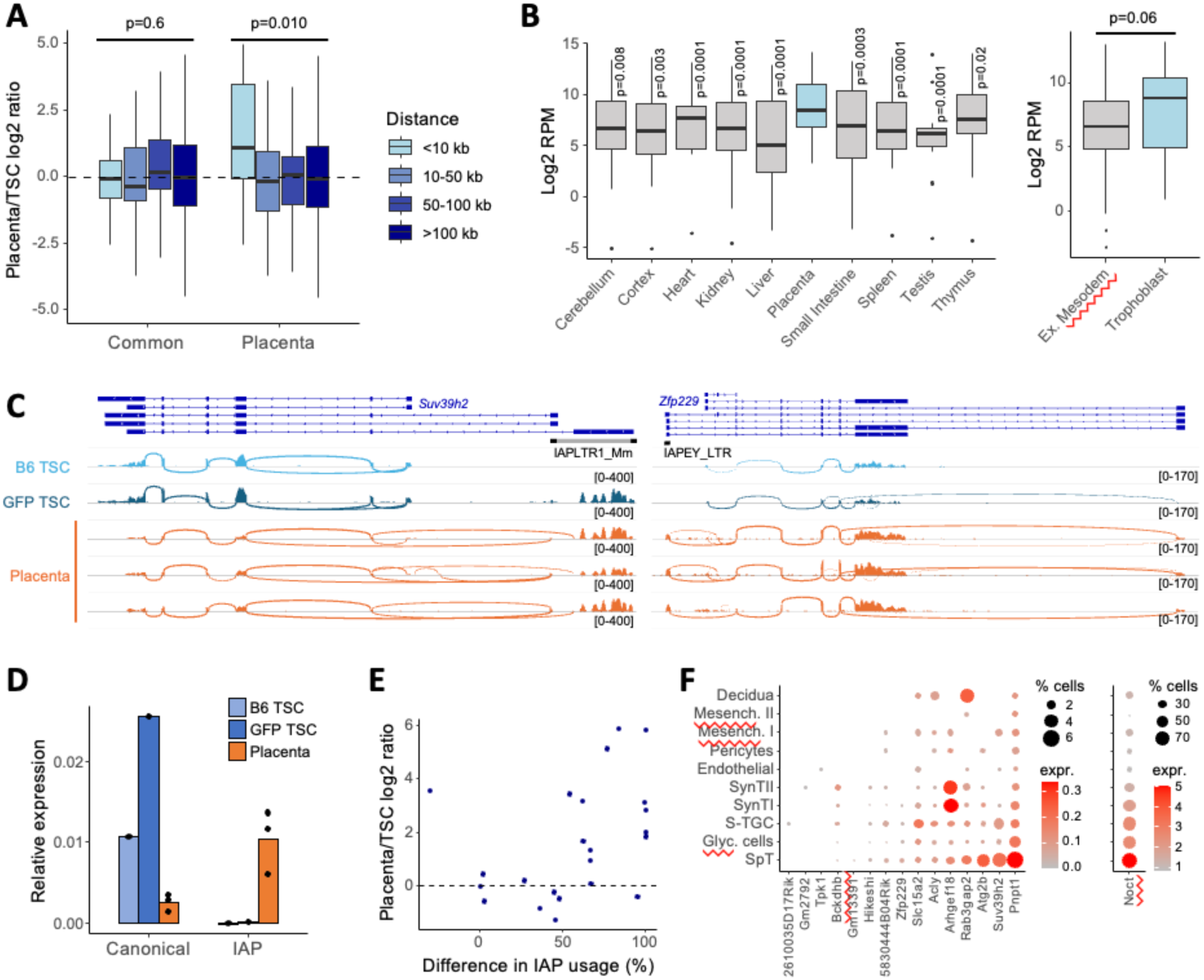
IAP elements generate placenta-specific TSSs. A) Relative expression (Placenta/TSCs) of genes containing TSSs within the indicated distances from lAPs with H3K4me3 in TSCs and placenta (common) or only in the placenta. B) Expression of genes within 10 kb of placenta-specific lAPs in ENCODE or our own data. C) Examples of genes with placenta-specific lAP-derived TSSs. D) RT-qPCR data for the canonical or lAP-derived isoforms of *Suv39h2.* E) Relative expression (Placenta/TSCs) of genes with lAP-derived isoforms as a function of the difference in the percentage of total transcription that is initiated at lAPs. F) Expression of genes with lAP-derived isoforms in placenta snRNA-seq data.

Several genes displayed a correlation between switching to an IAP-derived TSS and increased expression in placenta (relative to TSCs), but this association was not seen for all genes (Figure 3E), including *Suv39h2*. We hypothesised that isoform switching may instead (or additionally) drive gene expression within a specific placental compartment. Analysis of snRNA-seq showed that several IAP-associated genes (including *Suv39h2*) have prominent expression in spongiotrophoblast, glycogen cells and/or sinusoidal trophoblast giant cells (Figure 3F), in line with the overall IAP expression patterns (Figure 2F).

These data demonstrate that activation of young TEs (especially IAPs) in mouse placenta can drive cell-specific expression of genes with known roles in placental development.

### Epigenetic silencing of IAPs affects expression of nearby genes

To test the extent by which IAPs shape trophoblast gene expression, we performed CRISPR interference (CRISPRi) on TSCs. We first established a TSC line carrying a dCas9-KRAB transgene, then introduced a sgRNA targeting young IAPEz elements belonging to the IAPLTR1 and IAPLTR2 families (Figure 4A,B). Analysis of TE subfamily expression from CRISPRi RNA-seq data revealed a moderate downregulation of IAPLTR1_Mm elements, which were the main IAP subfamily expressed in our transgenic TSC line (Figure S6A). Because subfamily-level analyses can be confounded by signal from TE fragments within larger transcriptional units, we also evaluated the expression of individual IAP elements (using uniquely mapped reads), which showed clear downregulation of highly expressed loci (Figure 4C). Despite the relatively moderate silencing of IAP elements by CRISPRi, we found that genes within 10kb of IAPLTR1_Mm elements displayed reduced expression when compared to genes close to elements from other IAP subfamilies (Figure 4D). We also observed a similar but more subtle effect when analysing all IAPs predicted to be targeted by the sgRNA (Figure S6B), which include non-expressed elements.

**Figure 4.**
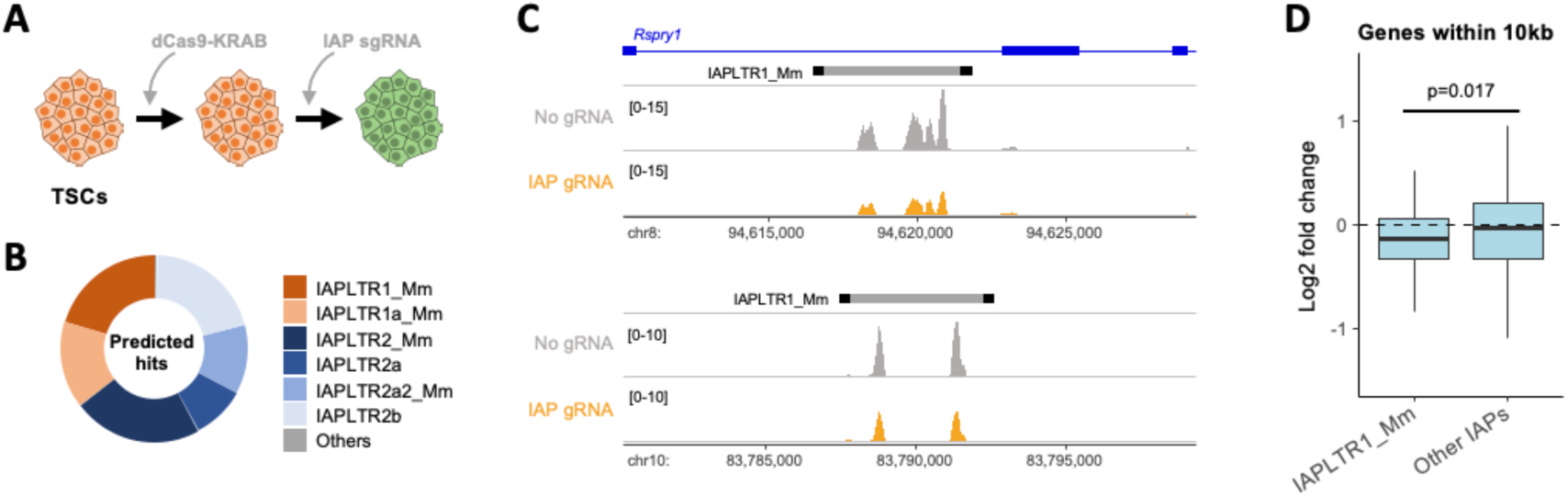
IAP silencing affects expression of nearby genes. A) Schematic of the CRISPRi experiment. A stable TSC line was established using PiggyBac dCas9-KRAB and lentiviral sgRNA-GFP constructs. B) In silico prediction of the IAP elements targeted by the IAP sgRNA. C) RNA-seq profiles (unique reads) at two expressed IAP loci in control or IAP sgRNA CRISPRi TSCs. D) Relative expression (IAP sgRNA over control) of genes within 10 kb of elements of the IAPLTR1_mM subfamily or other IAP subfamilies.

Notably, differential gene expression analysis revealed pronounced transcriptional differences in IAP CRISPRi TSCs (Figure S6C; Table S4). Upregulated genes (n=122) were related to cell migration and adhesion (Figure S6D), although there were no obvious changes in cell morphology. Downregulated genes (n=164) were linked to vascular development (Figure S6D), including genes such as *Nos3* (endothelial nitric oxide synthase) and *Eng* (endoglin), which regulate uterine artery remodelling and impact fetal growth^47,48^, and may act from trophoblast in a paracrine fashion. Whilst many of these gene expression differences are likely to be indirect downstream effects of IAP manipulation, they nonetheless suggest a potential for important phenotypic consequences during pregnancy. Notably, there are four downregulated genes (and no upregulated ones) within close proximity of an IAPLTR1_Mm element, including *Thbd* (thrombomodulin), the knockout of which is embryonically lethal due to defects in placental development^49,50^.

### TEs provide species-specific regulatory elements in mouse placenta

Previous work comparing the genomes of *Mus musculus*, *Mus caroli* (Ryukyu mouse, diverged from *musculus* 3 Mya), *Mus pahari* (Gairdner’s shrewmouse, diverged from *musculus* 6 Mya) and rat highlighted how recent TE activity shaped protein-coding genes and regulatory activity within the Muridae lineage^51^. We used these genomes to characterise the evolution of TE subfamilies with regulatory potential in mouse trophoblast. The RLTR13D5 subfamily, which has regulatory activity in mouse TSCs, is absent in rat (Figure 5A), as previously shown^11^. Out of the subfamilies identified here as having regulatory potential in the placenta, RMER10A has a high percentage of orthologous elements between *musculus* and the other species analysed (Figure 5A). In contrast, IAPs and young mouse L1s only expanded after the split between *caroli* and *musculus* (Figure 5A).

To evaluate whether TEs make a species-specific contribution to placental gene regulation, we collected placentas from *Mus musculus* and *Mus pahari*. Despite stage matching the embryos relative to *Mus musculus* E14.5, placentas looked dramatically different across the two species (Figure S7). The trophoblast of the *pahari* placentas appeared denser and more deeply implanted than *musculus*, and displayed a markedly reduced elaboration of the CD31-positive vascular endothelial tree. This resulted in reduced intercalation of the CK7-positive trophoblasts with the vascular network that is reminiscent of an earlier stage musculus placenta^52^ (Figure S7). Similarly in the junctional zone, the differentiation of mature spongiotrophoblasts from NCAM1-positive junctional zone progenitors^23^ appeared to be delayed in the pahari placenta (Figure S7B). RNA-seq uncovered 3,884 differentially expressed genes between the placentas of both species (Table S5). Notably, these included higher expression of markers of labyrinth and junctional zone progenitors in *pahari* placenta, with concomitant lower expression of markers of their differentiated counterparts (Figure S8A), which is congruent with the histological observations. To ask whether the placenta diverged more than other organs between the two species, we also performed RNA-seq on a small collection of *pahari* adult tissues (brain, kidney, liver and spleen) and compared them with *musculus* data from ENCODE. These data suggested that the placental transcriptome is more distinct between the two species when compared to other tissues (Figure S8B), which is in line with the faster evolutionary diversification of this unique organ.

We then performed CUT&Tag on *pahari* tissues and tested for enrichment of TE subfamilies overlapping peaks of histone modifications, as we previously did for *musculus* tissues. Similar to *musculus*, *pahari* placenta did not stand out amongst other tissues regarding the association of TE subfamilies with regulatory marks, but notably RMER10A was also enriched in this species (Figure S8C, Table S6). We then repeated the same analysis focusing on orthologous regions, and separating them based on whether CUT&Tag peaks are present in both species (epigenetically conserved) or only in one (epigenetically divergent). Unexpectedly, this analysis revealed that more TE subfamilies were enriched for epigenetically divergent peaks (including RMER10A) than conserved ones (Figure S8D,E, Table S7,S8). This indicates that distinct subsets of RMER10A elements bear active chromatin marks between the two species. Indeed, read coverage at orthologous RMER10A elements displayed little correlation between the two species (Figure 5B), as also exemplified by specific loci (Figure 5C). The species-specific activity of orthologous RMER10A elements could be due to genetic divergence and/or differences in expression of relevant transcription factors, amongst other possibilities. Finally, we also performed TE subfamily enrichment analysis on non-orthologous CUT&Tag peaks, which showed a pronounced enrichment for some IAP and L1 subfamilies (Figure 5D, Table S7,S8), suggesting that these TEs make a significant contribution to the species-specific regulatory landscape in the *musculus* placenta.

**Figure 5.**
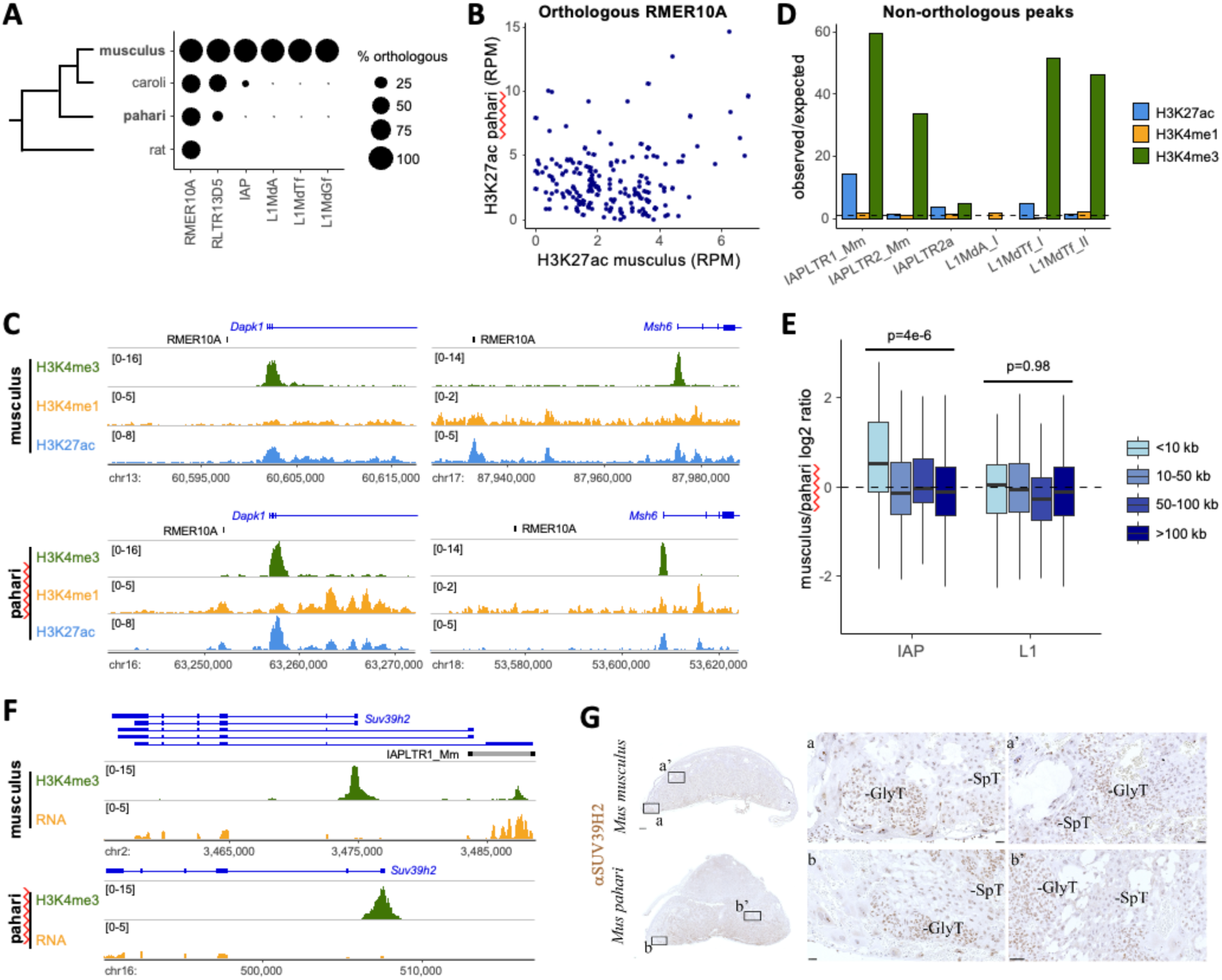
Comparative analysis of TE-mediated regulation in. *Mus* **species. A)** Percentage of *Mus* musculus TEs from the indicated subfamilies that display orthology in *Mus caroli, Musoahariand* rat. B) Placenta H3K27ac CUT&Tag signal at RMER10A elements orthoiogous between *musculus* and pahan (only elements with an H3K27ac peak in at least one species are plotted). C) Examples of loci with epigenetically divergent orthoiogous RMER10A elements between *musculus* and pahan placentas. D) Enrichment of the indicated TE subfamilies for muscu/us-specific histone modification peaks. E) Relative placental expression *(musculus/pahari)* of genes containing TSSs within the indicated distances from H3K4me3-marked IAP or L1 elements. F) H3K4me3 and RNA profiles for *Suv39h2* in *musculus* and pahan placenta. G) SUV39H2 immunostaining of sagjttaHy-sectioned placentas at the midline for *Mus musculus* and *Mus pahari.* Left: whole placenta is oriented with the maternal decidua at the top and fetal chorionic plate at the bottom, Positive staining is brown, counterstained with haemotgxylin (blue). Clusters of glycogen trophoblast cells (GlyT) are indicated, alongside junctional zone spongiotrophoblasts (SdT). Scale bars show 250 pm (whole placenta *Mus musculus),* 50 pm (whole placenta *Mus pahari),* 25 pm (insets a, a’ and b) and 50 pm (inset b’).

To test for potential effects of TEs on species-specific gene expression in the placenta, we compared the expression levels of genes proximal to epigenetically active TEs between *musculus* and *pahari*. For RMER10A elements, we found only a weak bias in expression for genes close to elements with species-specific H3K27ac peaks (Figure S8F). In contrast, genes within 10 kb of H3K4me3-marked IAP elements showed a clear preference for expression in *musculus* placenta (Figure 5E, S8G), supporting the hypothesis that IAP insertions drive species-specific gene expression differences. No such bias was seen with genes proximal to H3K4me3-marked L1s (Figure 5E). Out of 29 genes within 10 kb of an active IAP element, 21 had higher expression in *musculus* placenta relative to *pahari* (Table S9), which included the chimeric transcript driving *Suv39h2* expression in *musculus* placenta (Figure 5F). We also asked whether the presence of this IAP-driven *Suv39h2* transcript in *musculus* led to species-specific patterns of SUV39H2 protein expression. However, immunohistochemistry revealed largely similar expression patterns in *musculus* and *pahari* placentas, with strong signal in trophoblast cells of the labyrinth layer and in glycogen cells of the junctional zone, but reduced expression in mature spongiotrophoblast (Figure 5G). There appears to be higher SUV39H2 expression in *musculus* versus *pahari* spongiotrophoblast, which would be congruent with species-specific IAP expression in this cell type, but confirmation would require more quantitative approaches.

These comparative analyses support the hypothesis that the insertion of IAP elements close to genes can affect their expression in the placenta, with potential consequences for development and evolution.

## Discussion

We have queried the gene regulatory potential of TEs in the mouse placenta, building on work from TSCs that suggested a prominent role for ERVs as enhancers in mouse trophoblast^11,12^. Surprisingly, in contrast with human placenta, we found that the mature placenta contained almost no TE subfamilies enriched for canonical enhancer-associated epigenetic marks. Our results suggest that the mouse placenta is not exceptional with respect to the co-option of TEs as regulatory elements. Indeed, the increasing number of TE co-option examples seen in other tissues seems to challenge this common assumption^10,53^. A recurrent argument for why TE co-option might be more prevalent in the placenta is that this organ is particularly permissive for TE expression, given its relatively low levels of DNA methylation^4^. However, there is clear evidence that young LINE-1 elements^34^ and multiple ERV subfamilies^54^ remain under careful epigenetic control in mouse trophoblast. ERV deregulation in the mouse placenta has been associated with the onset of a deleterious inflammatory response via viral mimicry^54^, which could justify the need to maintain ERV silencing. The regulatory potential of TEs may therefore be just as epigenetically constrained in the placenta as in other somatic tissues, thus limiting the opportunities for TE co-option. The extent by which a tissue relies on TEs for gene regulation might instead be far more dependent on the chance matching between tissue-specific transcription factors and binding to motif-carrying TEs within a given species, at the right place in the genome and at the right time in evolution to confer a selective advantage. Whilst maternal-fetal conflicts may create selective pressures for TE co-option in the placenta, epigenetic mechanisms can still act as gatekeepers and dictate the frequency of co-option events in this organ.

We nonetheless found that young IAP elements can exert regulatory effects on nearby genes, namely through the creation of tissue-specific transcriptional start sites. It is also possible that a subset act as enhancers, similar to young L1 elements in mouse and human embryonic stem cells^40–43^. Recent findings also support a role for IAPs in regulating trophoblast gene expression, but in preimplantation development^36^. Additionally, given that we observe robust expression of IAPs, there are other potential consequences that we have not explored here. Many IAP elements retain capacity for retrotransposition^55^ and could therefore contribute to somatic mosaicism in the placenta, although the potential functional impact of this stochastic action might be questionable. It is also possible that IAPs trigger and/or enhance antiviral responses within the placenta, similar to what has been suggested for other TEs^54,56^. Finally, several IAP elements can produce functional viral proteins, as well as viral-like particles^35,55^. IAP-derived Gag protein can be detected within the junctional zone of the placenta^34^, which could affect cellular processes, e.g., by binding to RNAs, similar to the Gag-like PEG10 protein^6,57^. Moreover, IAP-derived viral-like particles are found in the placenta and may cross into the maternal decidua^35^, creating an opportunity for maternal-fetal communication akin to the inter-neuronal RNA transfer mediated by Arc^58^.

IAPs have previously been proposed to play regulatory roles in the placenta^20^. Namely, an IAP insertion in the CF1 mouse strain controls the expression of the *Mipp* gene^59^, but this insertion is absent in the B6 strain used here. There are also multiple IAP elements lying close to pregnancy-specific glycoprotein genes^20,60^, which are highly expressed in the placenta. However, in our dataset we find no evidence of active chromatin marks at these IAPs, making their functional relevance unclear. Instead, we found several other examples of IAP elements acting as alternative promoters to genes with known roles in trophoblast development, including *Suv39h2*^44^. The impact of these IAP-encoded promoters on phenotypes remains to be explored. The potential to leverage TSCs for this purpose is limited, not only because this would only reveal cellular phenotypes, but also because the majority of IAP-derived promoters are only expressed in the mature placenta (and not detected in TSCs differentiated in vitro). Genetic and/or epigenetic manipulation of IAPs in vivo would be required to fully test their potential effect on placental and embryonic development. Such experiments may also help to establish links to inter-species and/or inter-strain differences in placental development, such as those seen here between *Mus musculus* and *Mus pahari*. For example, one hypothesis is that differences in *Suv39h2* expression imparted by the nearby IAP may affect trophoblast differentiation and thereby partly explain the seemingly delayed maturation of the *pahari* placenta.

IAPs are highly polymorphic across mouse strains^61^, which is reflective of their young evolutionary age. We confirmed that IAP elements identified here as active in the placenta also display high inter-strain genetic variation (Figure S9), suggesting that any one insertion is unlikely to play a critical role in development and may instead generate more subtle phenotypic variation. In addition to genetic polymorphisms, we showed that IAPs also display inter-individual variation in DNA methylation in the placenta (Figure S3E), similar to what has been observed in other tissues^38^. Inter-strain differences in KRAB zinc finger proteins impart further variation in IAP methylation across mouse strains^62^. Altogether, this suggests that a high degree of inter-strain and even inter-individual variation might exist in the regulatory effects of IAPs. The resulting phenotypic variation could provide opportunities for natural selection that would lead to the eventual fixation of IAPs that confer an adaptive advantage. The current snapshot of IAP genetics and activity may therefore be reflective of the early stages of co-option. Similar arguments have been made for IAP activity in the neuronal lineage, where a subset of elements escape TRIM28-mediated repression and affect the expression of nearby genes^63^. Given that IAPs also pose a high mutagenic risk, this constitutes an interesting model to understand how host genomes balance the need to maintain genome integrity with the potential benefits of harnessing the regulatory potential of TEs.

## Materials and Methods

### Mouse lines and tissue collection

All experimental procedures were performed under licenses by the Home Office (UK) in accordance with the Animals (Scientific Procedures) Act 1986. Mating of *Mus musculus* (C57BL/6) females was timed based on the appearance of a vaginal plug, and culling performed at E3.5 for blastocyst flushing or at E14.5 for placenta collection. As vaginal plugs are not formed in *Mus pahari*, females were culled when visibly pregnant, and placentas collected from conceptuses that were stage-matched to *musculus* E14.5 based on embryo morphology. Adult animals were used for the collection of brain, kidney, liver and spleen samples. To generate trophoblast and extraembryonic mesoderm samples for RNA-seq, ATAC-seq and Cut&Tag experiments, heterozygotes Meox2-cre females (Meox2^tm1(cre)Sor/+^)^64^ were crossed to homozygotes mTmG dual reporter line males (Gt(ROSA)26Sor^tm4(ACTB-tdTomato,-EGFP)Luo/J^)^65^. Mating was timed based on the presence of a vaginal plug and placenta were collected at E13.5. For the analysis of variably methylated IAPs, tails and whole placentas of C57BL/6J embryos were dissected at E16.5 and flash frozen in liquid nitrogen.

### Fluorescence Activated Cell Sorting (FACS)

FACS was performed as described in Vignola et al^22^. Briefly, placentas were examined under a microscope (Zeiss Axioplan 2) for GFP and Tomato fluorescence expression, and individual positive samples were sliced into small pieces using sterile scalpels and digested at 37°C for 1 hour and 45 minutes in a collagenase solution containing 0.1% Collagenase P (Roche-LifeScience, 11213865001), 0.1% BSA (Merck, A9647) in PBS, and 50ml DNase I (STEMCELL Technologies UK, 07900). During the digestion period, samples were mechanically dissociated every 15-30 minute interval, by gently pipetting up and down the solution firstly with a Pasteur pipette and after with needles of decreasing gauge size (18G, 21G and 23G). Samples were filtered through a 70mm cell strainer and centrifuged at 4°C for 5 minutes at 1200 g. Red blood cells were removed by resuspending the samples in RBC lysis buffer (BioLegend, 420301) for 5 minutes on ice with occasional shaking. Samples were washed once with PBS and cell pellets resuspended in 200ml of cell staining buffer (BioLegend, 420201) containing 4′,6-diamidino-2-phenylindole (DAPI, 1:5000). Cell sorting was performed using a FACSAriaTM III instrument as described in Vignola et al^22^. For the RNA-seq experiment, sorted cells were collected directly into TRIzol LS reagent (Thermo Fischer Scientific, 10296028). For the ATAC-seq and Cut&Tag experiments, 2-3 placentas within the same litter positive for the same fluorescent protein (GFP or Tomato) were pooled together and sorted cells were collected in 1X PBS and immediately subjected to ATAC-seq and Cut&Tag procedures.

### TSC derivation and culture

Conditioned medium for the culture of TSCs (TS-CM) was prepared by incubating TS base medium (TS-BM; RMPI 1640 supplemented with 20% FBS, 1X antibiotic-antimycotic, 1 mM sodium pyruvate and 50 µM 2-mercaptoethanol) on confluent, mitomycin-treated primary mouse embryonic fibroblast (PMEF) feeders for 72 h, followed by filtration through a 0.45 µm PES membrane. Blastocysts from C57BL/6 mice were plated individually onto gelatin-coated wells containing mitomycin-treated PMEF feeders in medium consisting of 70% TS-CM and 30% TS-BM, supplemented with 50 ng/ml bFGF, 2 μg/ml heparin and 100 µM Y-27632 (ROCK inhibitor). Following hatching, once 3-4 layers of TSCs formed around the blastocyst, outgrowths were dissociated using 0.5% trypsin-EDTA. TSCs were passaged at sub-confluency until feeder cells were eliminated and subsequently cryopreserved in liquid nitrogen in freezing medium containing 90% culture medium and 10% DMSO. For regular passaging of established lines (B6 TSCs, GFP TSCs^21^ and Rs26 TSCs), medium with no ROCK inhibitor and reduced bFGF (25 ng/ml) and heparin (1 µg/ml) was used. TSC differentiation was performed by culturing cells in TS base medium for 4 days.

### Immunohistochemistry and immunofluorescence

Immunohistochemistry (IHC) was performed by incubating paraffin embedded sections for 20 minutes at 95°C in either Tris-EDTA pH 9 buffer (10mM Tris Base, 1mM EDTA) or 10 mM tri-sodium citrate pH 6 buffer, depending on the primary antibody utilised (Table S10). Slides were blocked in Casein buffer (Thermo Scientific, 37528) for 1 hour at room temperature. Protein detection was performed by incubating the sections with primary antibodies overnight at 4°C, followed by 1 hour incubation at room temperature with secondary antibodies. All antibodies are indicated in Tabel S10. Sections were incubated in ABC Staining solution (2BScientific, PK-6100) and stained using the DAB Peroxidase Substrate kit (2BScientific, SK-4100). Slides were counterstained with Haematoxylin (Sigma-aldrich, 51275) and mounted using DPX mounting medium (Merck Life Science Limited, 06522). Slides were scanned on a Hamamatsu Scanner 2.0-HT and pictured acquired using the NDP.view2 software. Immunofluorescence was performed by incubating the sections in Tris-EDTA pH 9 buffer as described above. Slides were blocked in Casein buffer, and proteins were detected by incubating the sections with primary antibodies overnight at 4°C, followed by 1 hour incubation at room temperature with fluorescence conjugated secondary antibodies. All antibodies are listed in Tabel S10. The Ready Probes Tissue Autofluorescence Quenching kit (Thermo Scientific, R37630) was used for autofluorescence quenching. The slides were mounted using Vectashield Antifade Mounting Medium (2BScientific, H-1700-10) and pictures acquired using a Zeiss Axio Observer Z1 microscope.

### CRISPRi

Rs26 TSCs were seeded at a density of 2.2 × 10^6^ cells per gelatin-coated 10 cm dish. Twenty-four hours later, cells were transfected using FuGENE 6 (Promega) according to the manufacturer’s instructions, with a transfection mix containing 600 µl Opti-MEM, 60 µl FuGENE 6, 2 µg Super PiggyBac Transposase plasmid (System Biosciences) and 3 µg PiggyBac dCas9-KRAB plasmid (Addgene #191314, a kind gift from the Wysocka laboratory). Cell selection was perfomed by FACS after induction of dCas9-KRAB-BFP expression with 8 µg/ml doxycycline hydrochloride for 72 hours. Stably transfected Rs26 KRAB TSCs were subsequently transduced with Lenti_sgRNA_EFS_GFP (Addgene #65656) carrying a single guide RNA targeting IAP retrotransposons (5’-GATGGGCTGCAGCCAATCA-3’) at a multiplicity of infection (MOI) of 10. Double-positive (BFP⁺/GFP⁺) populations were isolated by FACS. Rs26 KRAB sgIAP TSCs were then treated with 8 µg/ml doxycycline hydrochloride to induce CRISPR interference. Cells expressing dCas9-KRAB (treated with doxycycline hydrochloride) but lacking the sgRNA were used as controls, as we found that uninduced cells already expressed high levels of dCas9-KRAB. Gene expression changes were assessed by RT-qPCR 72 hours post-induction.

### CUT&Tag

CUT&Tag was performed using either 50,000 cells or 5 mg of tissue per antibody. Cells were harvested fresh using TrypLE Express and centrifuged at 600 × g for 5 min at room temperature (RT). Placentas were homogenised in 1 ml wash buffer (20 mM HEPES pH 7.5, 150 mM NaCl, 0.5 mM spermidine, 1X protease inhibitor cocktail (PIC), 0.1% BSA) with a Dounce homogeniser using a tight pestle 30 times. Dounced tissue was transferred to a 1.5 ml tube and centrifuged at 600 × g for 5 min at RT. Pellet was resuspended in 500 µl ice-cold nuclei extraction buffer (20 mM HEPES pH 7.5, 10 mM KCl, 0.5 mM spermidine, 1% Triton X-100, 20% glycerol, 1X PIC) and incubated on ice for 10 minutes. Nuclei were pelleted by centrifugation at 1,300 × g for 5 min at 4 °C. Pelleted cells or nuclei were washed twice with 1 ml wash buffer (20 mM HEPES pH 7.5, 150 mM NaCl, 0.5 mM spermidine, 1X PIC) by gentle pipetting. BioMagPlus Concanavalin A-coated magnetic beads (Generon) were activated by washing and resuspension in binding buffer (20 mM HEPES pH 7.5, 10 mM KCl, 1 mM CaCl_₂_, 1 mM MnCl_₂_). Activated beads (10 µL per sample) were added to the cells and incubated for 10 min at RT. Bead-bound cells were isolated using a magnetic stand, the supernatant was removed, and cells were resuspended in 100 µl antibody buffer (20 mM HEPES pH 7.5, 150 mM NaCl, 0.5 mM spermidine, 1X PIC, 0.05% digitonin, 2 mM EDTA, 0.1% BSA). Primary antibodies were added at a 1:10-1:100 dilution, and samples were incubated overnight at 4 °C on a nutator. Primary antibodies (Table S10) were added at a 1:10-1:100 dilution, and samples were incubated overnight at 4 °C on a nutator. Cells were incubated with secondary antibody diluted 1:100 in 100 µl Dig-Wash buffer (20 mM

HEPES pH 7.5, 150 mM NaCl, 0.5 mM spermidine, 1X PIC, 0.05% digitonin) for 1 h at RT on a nutator. Cells were incubated with secondary antibody (Table S10) diluted 1:100 in 100 µl Dig-Wash buffer (20 mM HEPES pH 7.5, 150 mM NaCl, 0.5 mM spermidine, 1X PIC, 0.05% digitonin) for 1 h at RT on a nutator. Cells were washed three times with 800 µl Dig-Wash buffer before incubation with Protein A-Tn5 transposase fusion protein (pA-Tn5), pre-loaded with Illumina Nextera adapters (kind gift from the Henikoff laboratory via the Madapura laboratory). pA-Tn5 was diluted 1:250 in Dig-300 buffer (20 mM HEPES pH 7.5, 300 mM NaCl, 0.5 mM spermidine, 1X PIC, 0.05% digitonin), and 100 µl was added per sample followed by incubation for 1 h at RT on a nutator. Unbound pA-Tn5 was removed by three washes with 800 µL Dig-300 buffer. Cells were then resuspended in 300 µl tagmentation buffer (10 mM MgCl_₂_ in Dig-300 buffer) and incubated at 37 °C for 1 h on a nutator. Tagmentation was stopped by the addition of 10 µl 0.5 M EDTA, 3 µl 10% SDS, and 2.5 µl Proteinase K (20 mg/ml), followed by overnight incubation at 37 °C. DNA was extracted using phenol:chloroform:isoamyl alcohol (25:24:1, v/v; Sigma-Aldrich) and Phase Lock Gel Heavy tubes (5 PRIME), followed by chloroform extraction and ethanol precipitation. DNA pellets were washed with 100% ethanol, air-dried, and resuspended in 30 µl TE buffer (10 mM Tris-HCl pH 8.0, 1 mM EDTA) containing RNase A (1:400). Libraries were amplified using 8 µl DNA per sample with NEBNext HiFi 2X PCR Master Mix, universal i5 primer, and uniquely indexed i7 primers (0.5 µM each). PCR conditions were 72 °C for 5 min, 98 °C for 30 s, followed by 12 cycles of 98 °C for 10 s and 63 °C for 10 s, with a final extension at 72 °C for 1 min. Libraries were purified using AMPure XP beads (Beckman Coulter) according to the manufacturer’s instructions.

### ATAC-seq

ATAC-seq was performed as previously described^66^. Cells were centrifuged at 500 × g for 5 min at 4 °C. Pellet was resuspended in 50 µl ice-cold resuspension buffer (10 mM Tris-HCl pH 7.4, 10 mM NaCl, 3 mM MgCl_₂_, 0.1% Tween-20). Nuclei were centrifuged at 500 × g for 10 min at 4 °C. Pellet was resuspended in 50 µl transposition buffer (10 mM Tris-HCl pH 7.6, 5 mM MgCl_₂_, 10% dimethyl formamide, 100 nM transposase, 0.01% digitonin, 0.1% Tween-20, 33% PBS) and incubated at 37 °C for 30 min in a thermomixer set to 1000 rpm. DNA was eluted in 20 µl elution buffer using a commercial DNA purification kit (Zymo Research). Libraries were amplified using 20 µl DNA with NEBNext HiFi 2X PCR Master Mix, universal i5 primer, and uniquely indexed i7 primers (1.25 µM each). PCR conditions were 72 °C for 5 min, 98 °C for 30 s, followed by 5 cycles of 98 °C for 10 s, 63 °C for 30 s and 72 °C for 1 min. Libraries were eluted in 20 µl elution buffer using the same DNA purification kit.

### RNA-seq and RT-qPCR

Total RNA was isolated using a commercial RNA extraction kit (Qiagen) and treated with DNase I (Ambion) to remove residual genomic DNA. For RNA-seq library preparation, 500 ng of total RNA was subjected to ribosomal RNA depletion using the NEBNext rRNA Depletion Kit v2. Strand-specific libraries were generated using the NEBNext Ultra II Directional RNA Library Prep Kit for Illumina according to the manufacturer’s instructions. For RT-qPCR analysis of *Suv39h2* transcript isoforms, 1 µg of total RNA was reverse-transcribed using RevertAid Reverse Transcriptase (Thermo Scientific). Quantitative PCR was performed using KAPA SYBR FAST Master Mix (Sigma-Aldrich) and isoform-specific primers (Table S11) on a Roche LightCycler 480 system. Amplification was carried out for 40 cycles.

### Western Blotting

Cells were washed twice with ice-cold PBS and lysed in RIPA buffer (50 mM Tris-HCl pH 8.0, 150 mM NaCl, 1% NP-40, 0.5% sodium deoxycholate, 0.1% SDS) supplemented with protease inhibitor cocktail. Mature placenta tissue was homogenised in RIPA buffer using a mechanical homogeniser. Lysates were incubated on ice for 30 minutes with periodic agitation and clarified by centrifugation at 14,000 × g for 15 minutes at 4°C. Protein concentration was determined by BCA assay. Equal amounts of protein (10-30 µg) were denatured in Laemmli sample buffer containing β-mercaptoethanol at 95°C for 5 minutes and resolved by SDS-PAGE on 4-12% Bis-Tris polyacrylamide gels (Bolt, Invitrogen) using MES running buffer. Proteins were transferred onto nitrocellulose membranes using a semi-dry transfer system (Trans-Blot Turbo, Bio-Rad) according to the manufacturer’s instructions. Membranes were blocked in 5% BSA in PBST for 1 hour at room temperature and incubated with primary antibodies (Table S10) diluted in 5% BSA in PBST overnight at 4°C. Following three washes in PBST, membranes were incubated with HRP-conjugated secondary antibodies (Table S10) diluted in 5% BSA in PBST for 1 hour at room temperature. After three further washes in PBST, chemiluminescent signal was detected using Clarity Western ECL Substrate (Bio-Rad) and imaged on a ChemiDoc MP imaging system (Bio-Rad).

### Bisulphite pyrosequencing

Genomic DNA from tail and placenta samples was isolated by proteinase K digestion followed by phenol–chloroform extraction and ethanol precipitation. Bisulphite conversion was performed using the Imprint® DNA Modification Kit (Sigma-Aldrich) following the manufacturer’s two-step protocol. Pyrosequencing assays were designed with PyroMark Assay Design SW 2.0 (QIAGEN) and primer sequences are listed in Table S11. Bisulphite-converted DNA was PCR-amplified using biotinylated reverse primers (HotStarTaq; QIAGEN) under the following cycling conditions: 95°C for 3 min; 40 cycles of 94°C for 30 s, optimized annealing temperature for 30 s, 72°C for 55 s; final extension at 72°C for 5 min. Amplicons were processed on the PyroMark Q96 platform (Vacuum Workstation and Q96 MD pyrosequencer; QIAGEN) with PyroMark Gold Q96 reagents according to the manufacturer’s instructions. Percent methylation at each CpG was quantified using Pyro Q-CpG 1.0.9 (Biotage). Values were averaged across triplicate PCR reactions and then across CpGs for each locus.

### High-throughput sequencing data alignment

Sequencing reads were trimmed using Trim_galore, with default settings. Mapping was done with either Bowtie2^67^ (for CUT&Tag and ATAC-seq; default settings) or Hisat2^68^ (for RNA-seq; with --no-softclip) to the reference genome of the species of origin: mm10 for *Mus musculus* and GCF_900095145 for *Mus pahari*. Unless indicated, analyses were performed on reads with MAPQ of 2 or higher, which excludes reads with equal best mapping scores at multiple locations. TEtranscripts^32^ and SQuIRE^33^ were used to measure TE family-wide expression from RNA-seq data including multimapping reads.

### Peak detection and enrichment at repeat families

Ahead of peak detection, regions with abnormally high read density reads in the CUT&Tag IgG control were filtered out of BAM alignment files. ATAC-seq peak detection was performed using MACS2^69^ (with -q 0.05). For CUT&Tag peaks we used GoPeaks^70^ with the -r option set at the total number of reads divided by 10^6^, to take into account differences in sequencing depth; additionally, -t 200 was used for H3K4me3 and H3K27ac, and -t 2000 -l 200 for the broader H3K4me1 signal. Processed ChIP-seq peaks from multiple *Mus musculus* tissues were downloaded from ENCODE (Table S12). To measure enrichment at repeats, the number of ATAC-seq or CUT&Tag peaks overlapping each Repeatmasker-annotated repeat family (excluding tRNAs, simple and low complexity repeats) was compared with overlap frequencies across 1000 random controls (shuffled peaks, avoiding unmappable regions of the genome), yielding enrichment values and associated p-values (corrected for multiple comparisons). Significantly enriched repeat families had p<0.05, >2-fold enrichment, and at least 10 copies overlapped by peaks.

### TE and peak orthology

*Mus musculus* TE orthologues were identified in other rodent genomes using liftOver (with -minMatch=0.5), then filtering for elements that were annotated as belonging to the same subfamily in both species. LiftOver was also used to classify placental CUT&Tag peaks with regards to their orthology between *Mus musculus* and *Mus pahari*, as follows: 1) peaks that did not lift over were classified as non-orthologous, 2) peaks that lifted over and overlapped with a peak in both species were classified as epigenetically conserved, 3) peaks that lifted over but overlapped a peak in only one of the species were classified as epigenetically divergent.

### RNA-seq analysis

Strand-specific RNA-seq gene/TE raw counts or RPM values were extracted using the RNA-seq pipeline in Seqmonk. For measuring expression of individual IAP and RLTR4 elements, annotations of merged LTRs and internal fragments were generated using the “One code to find them all” tool^71^, keeping only intergenic elements for further analysis. For comparative analyses, *Mus musculus* and *Mus pahari* RNA-seq data were merged based on one-to-one gene orthologues. DESeq2^72^ was used to perform differential expression analysis, with padj<0.05 and 2-fold change thresholds. Gene ontology analysis was performed using topGO. To evaluate putative enhancer effects, absolute or relative gene expression levels were determined and grouped based on the distance between gene transcriptional start sites and the nearest H3K27ac-marked TE from each relevant subfamily. TE-gene chimeric transcripts were identified by performing transcriptome assembly on TSC and placenta data using Stringtie^73^ v2.2.3 (with --rf and guided by the Gencode vM25 annotation), then filtering for transcripts with transcriptional start sites overlapping TEs and with at least one exon overlapping Gencode-annotated exons. The list of IAP- and L1-derived transcripts was manually curated by browsing annotations and underlying RNA-seq data, which mostly removed transcripts where the annotated Gencode gene was only the TE itself. Quantification of transcript abundance was performed on a merged annotation of placental transcripts using Stringtie. Sashimi plots were generated using the IGV genome browser.

### Single-nuclei/cell RNA-seq analysis

The E14.5 mouse placenta single-nuclei dataset from Marsh et al.^23^ was used throughout. CIBERSORT^74^ was used to perform cell type deconvolution of bulk RNA-seq datasets, using the Marsh et al.^23^ dataset to create a placenta-specific gene signature matrix. For gene and IAP expression analysis, Cell Ranger was used with a custom annotation merging Gencode vM25 and a list of intergenic proviral IAP elements (LTRs and internal fragments merged as described above) expressed in either TSCs or placenta. Cell type clusters were annotated based on markers used in Marsh et al^23^.

### Estimating TE *cis*-regulatory activities

TE *cis*-regulatory activities were estimated by modelling changes in gene expression between two conditions as a linear combination of integrant occurences located in the vicinity of protein coding gene promoters^24^. A mouse version of the *cis*-regulatory susceptibility matrix *N* was generated, following the methodology previously described^24^. Assuming conservation of the *cis*-regulatory window size between humans and mice, integrant occurences were weighed using a gaussian kernel with standard deviation *L* set to 250,000 base pairs. Single-cell RNA-seq data from E6.5-E8.5 embryos^25^ were used to predict TE *cis*-regulatory activity in early trophoblast development, and snRNA-seq data from E14.5 placenta^23^ were used for differentiated trophoblast cell types. Pseudobulk datasets were generated for each cell type using the SingleCellExperiment R package and the cluster annotations generated in those studies. TE *cis*-regulatory activities were estimated using default parameters.

## Data and code availability

CUT&Tag, ATAC-seq and RNA-seq data generated for this study have been deposited in NCBI’s Gene Expression Omnibus under accession number GSE316744. Additional datasets had been previously deposited under the accession numbers GSE200761 (GFP TSC CUT&Tag) and GSE313753 (RNA-seq of sorted placental cells). Details of other external datasets used can be found in Table S12. All code associated with the manuscript is available via GitHub at https://github.com/MBrancoLab/Amante_2026_mPlacenta.

## Acknowledgements

We thank Jordi Lopez-Tremoleda and James Gillett for their efforts in setting up husbandry and experimental mating conditions for *Mus pahari* animals, Tania Maffucci for technical assistance with western blotting, and Didier Trono for scientific discussions.

