## Supplementary Figures for "IAP retrotransposons contribute to the transcriptional diversity of the murine placenta"

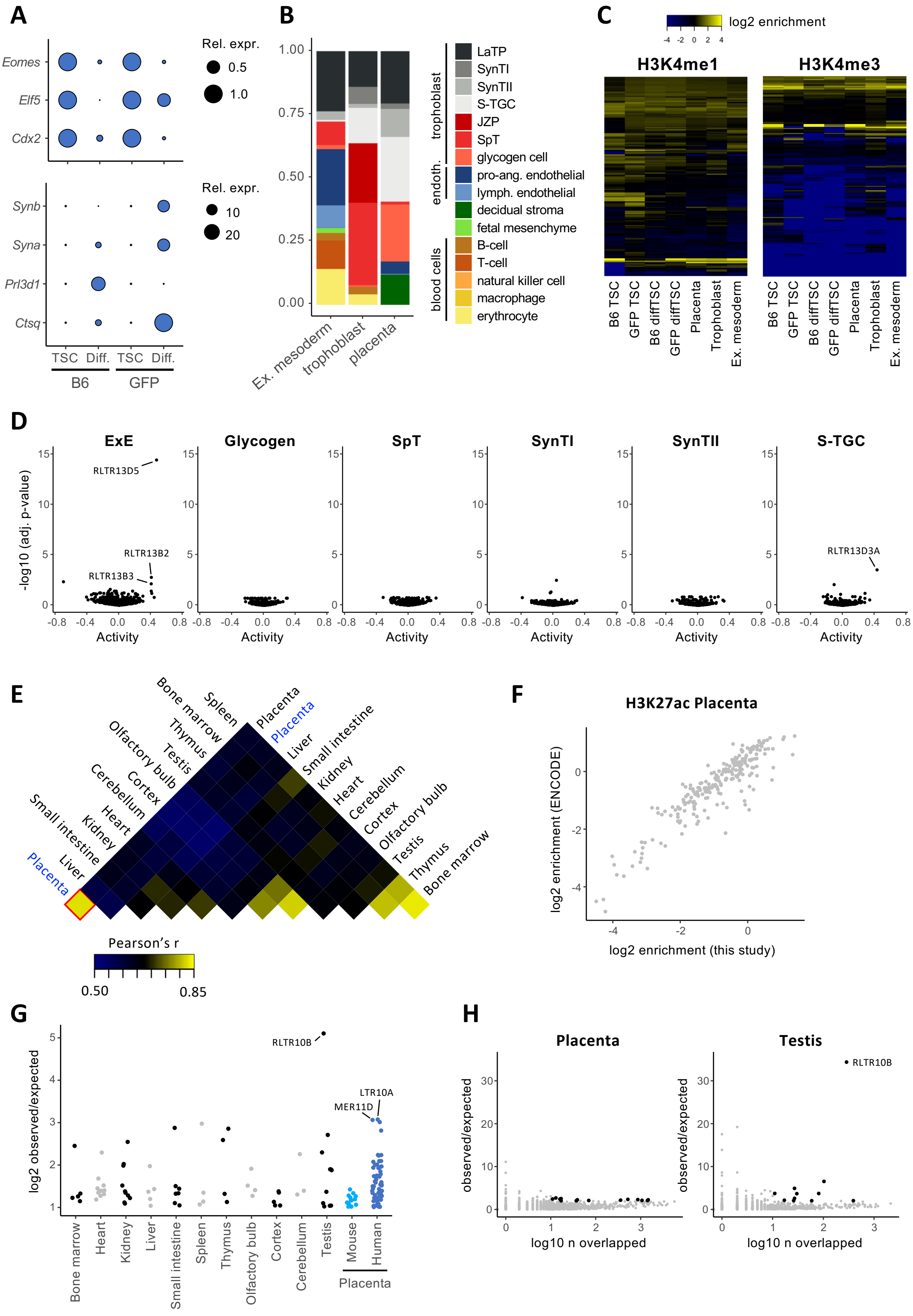


**Figure S1 – Regulatory TE landscape in mouse placenta.** A) Relative expression (RT-qPCR normalised to TSC) of markers of TSCs (*Cdx2*, *Elf5*, *Eomes*), syncytiotrophoblast (*Syna*, *Synb*) and trophoblast giant cells (*Prl3d1*, *Ctsq*) in B6 and GFP TSCs grown under stem cell or differentiation conditions. B) Cell-type deconvolution of bulk RNA-seq data using CIBERSORT and a placental snRNA-seq dataset. C) TE subfamilies displaying enrichment (observed/expected) for elements overlapping H3K4me1 or H3K4me3 peaks in at least one dataset. D) Prediction of TE regulatory activity from scRNA-seq data using craTEs. Epiblast was used as a reference for extraembryonic ectoderm (ExE), and endothelial cells as a reference for placental cell types (SpT – spongiotrophoblast, SynT – syncytiotrophoblast, S-TGC – sinusoidal trophoblast giant cells). E,F) Correlation of H3K27ac data between ENCODE (black) and our (blue) datasets. G) TE subfamilies enriched for H3K27ac peaks in the indicated tissues. H) TE subfamily H3K27ac enrichment in placenta or testis (black data points are classified as significantly enriched).

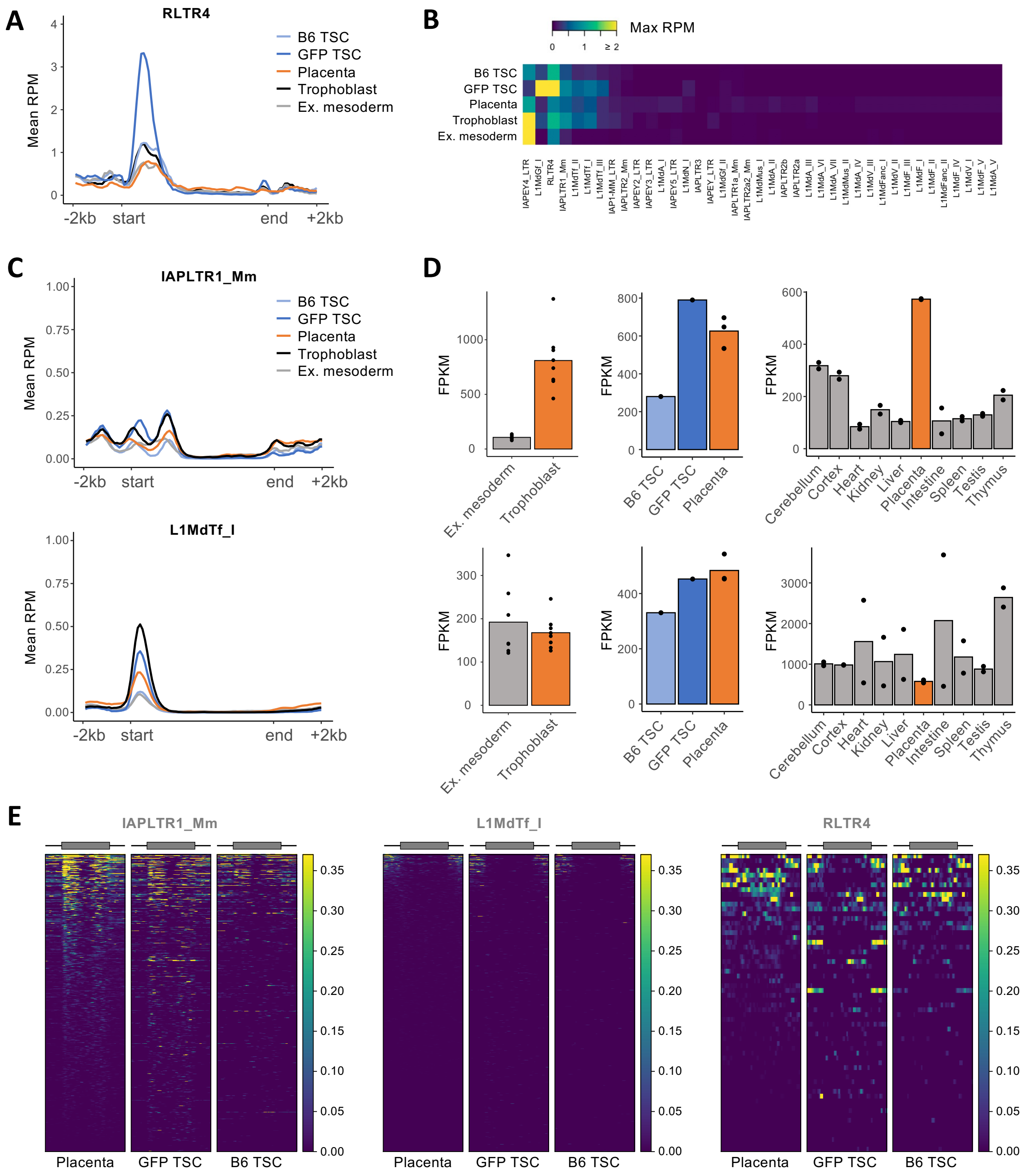


**Figure S2 – IAP and L1 expression.** A) Mean H3K4me3 CUT&Tag signal over RLTR4 elements, allowing for random assignment of non-unique reads. B) Maximum value in the mean H3K4me3 profile of the indicated TE subfamilies. C) Mean H3K4me3 CUT&Tag signal over IAPLTR1_Mm or L1MdTf_I elements, using only unique reads. D) Expression of IAPLTR1_Mm (top) or L1MdTf_I (bottom) subfamilies in our own or ENCODE datasets, with non-unique reads assigned using SQuIRE. E) RNA-seq signal (unique reads only) over the indicated TE subfamilies.

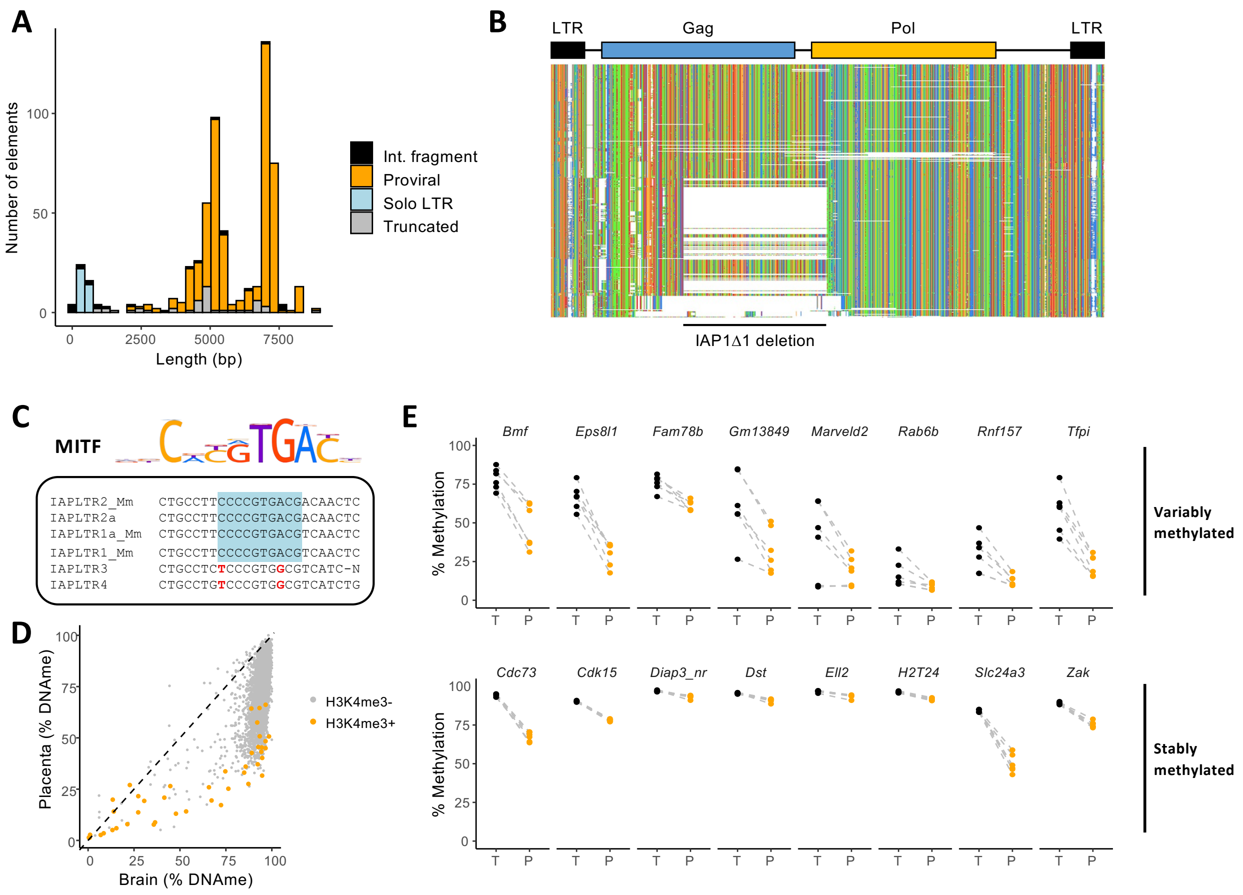


**Figure S3 – Characteristics of expressed IAPs.** A) Lengths of IAPs preferentially expressed in the placenta, subdivided by their genomic arrangement. B) Multiple alignment of proviral IAPLTR1_Mm and IAPLTR1a_Mm elements expressed in the placenta. C) MITF binding motif, which is present in IAPLTR1/2 elements (high placenta expression), but is mutated in IAPLTR3/4 elements (low/no placenta expression). D) DNA methylation levels of IAP LTRs in brain and placenta, with H3K4me3-marked elements in orange. E) DNA methylation levels of IAPs near the indicated genes in matched tail (T) and placental (P) samples from littermates. IAPs are classified based on their methylation variability in tail samples.

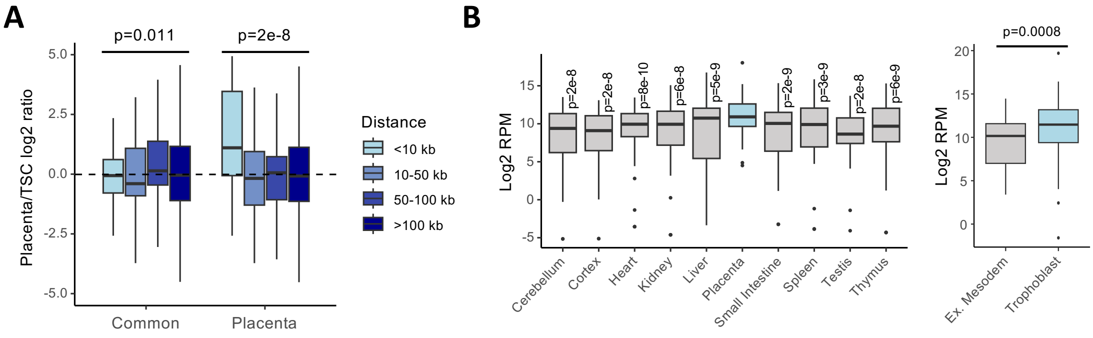


**Figure S4 – Expression of L1-proximal genes.** A) Relative expression (Placenta/TSCs) of genes containing TSSs within the indicated distances from L1s with H3K4me3 in TSCs and placenta (common) or only in the placenta. B) Expression of genes within 10 kb of placenta-specific L1s in ENCODE or our own data.

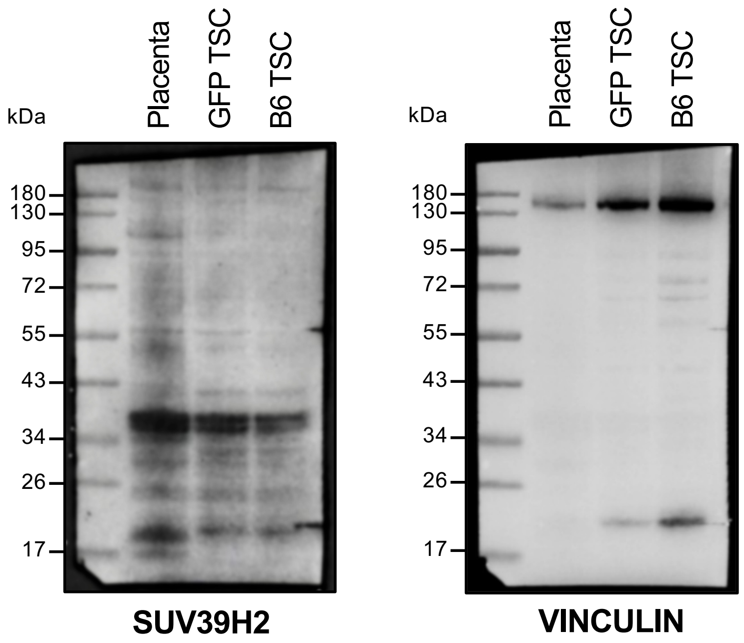


**Figure S5 – SUV39H2 protein profiles.** Western blot for SUV39H2 and VINCULIN (loading control) in E14.5 placenta, GFP TSC and B6 TSC. Whilst the IAP-driven *Suv39h2* transcript expressed in the placenta has the potential to produce a protein that is 28 kDa larger than the canonical form, there is no major shift in the main band detected when compared with TSCs. The anti-SUV39H2 monoclonal antibody used (Abcam ab190870) detects a C-terminal portion of the protein and has been previously validated in *Suv39h2*-depleted mouse cells (Jing et al, 2020, PMID: 32698678).

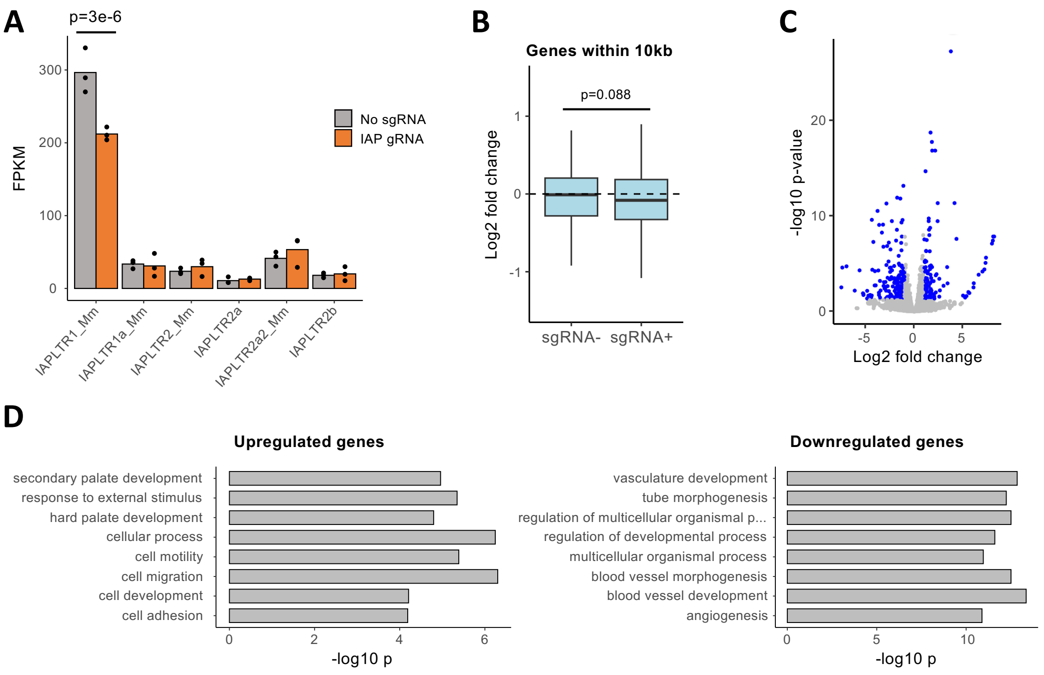


**Figure S6 – Transcriptional effects of IAP CRISPRi.** A) Subfamily-level expression of IAPs in CRISPRi, with non-unique reads assigned using SQuIRE. B) Relative expression (IAP sgRNA over control) of genes within 10 kb of elements of IAPs predicted to be targeted (or not) by the sgRNA. C) Volcano plot of gene expression changes after IAP CRISPRi, with differentially expressed genes highlighted in blue. D) Gene ontology analysis results of differentially expressed genes after IAP CRISPRi.

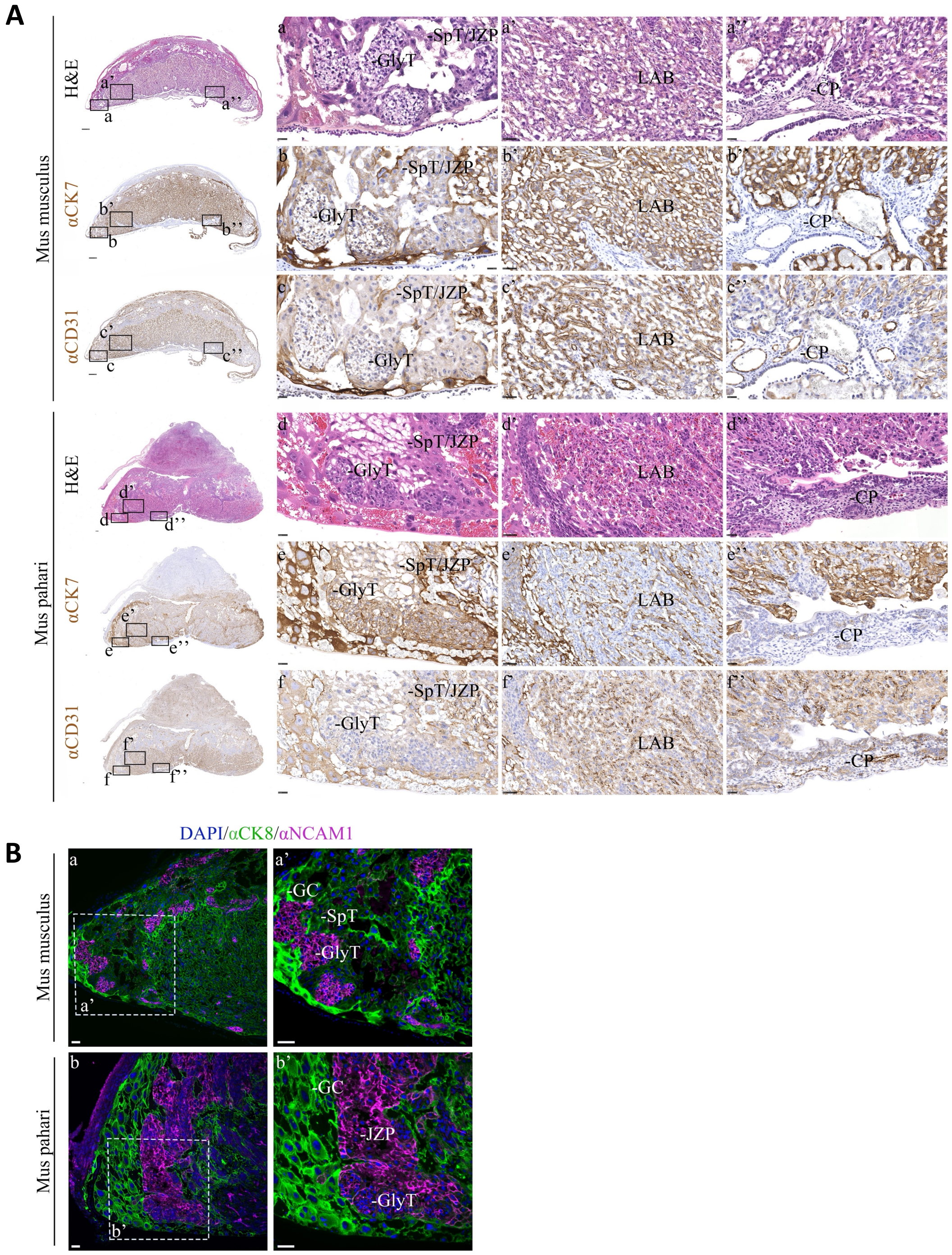


**Figure S7 – Comparative histological analysis of *Mus musculus* and *Mus pahari* placenta at E14.5.** A) Staining of sagittally-sectioned placentas at the midline for *Mus musculus* and *Mus pahari*. Left: whole placenta is oriented with the maternal decidua at the top and fetal chorionic plate at the bottom, stained with Haemotoxylin and Eosin (a, a', a'', d, d', d''), and immunohistochemistry for pan-trophoblast marker Cytokeratin 7 (CK7), (b, b', b'', e, e', e''), and the endothelial marker CD31 (c, c', c'', f, f', f''). Positive immunostaining is brown, counterstained with haemotoxylin (blue). Clusters of glycogen trophoblast cells (GlyT) are indicated, alongside junctional zone trophoblast that may be spongiotrophoblast or junctional zone progenitors (SpT/JZP). In the central labyrinth of the placenta (LAB), trophoblast cells are highly interdigitated with endothelial cells. The base of the placenta is defined by the chorionic plate (CP). Scale bars show 250 µm (whole placenta *Mus musculus*), 50 µm (whole placenta *Mus pahari*), 25 µm (insets a-f, a''-c'', f'') and 50 µm (insets a-f, a'-f'). B) Immunofluorescence detection of junctional zone trophoblast cells in *musculus* (a, a') and *pahari* (b, b') placentas at E14.5. All trophoblast cells are positive for cytokeratin 8 (CK8), with different levels of intensity between subtypes, e.g., Gly-T cells express low CK8, SpT cells are weakly positive while giant cells (GC) are strongly positive. NCAM1 is a marker of GlyT and JZP cells. Scale bars show 50 µm.

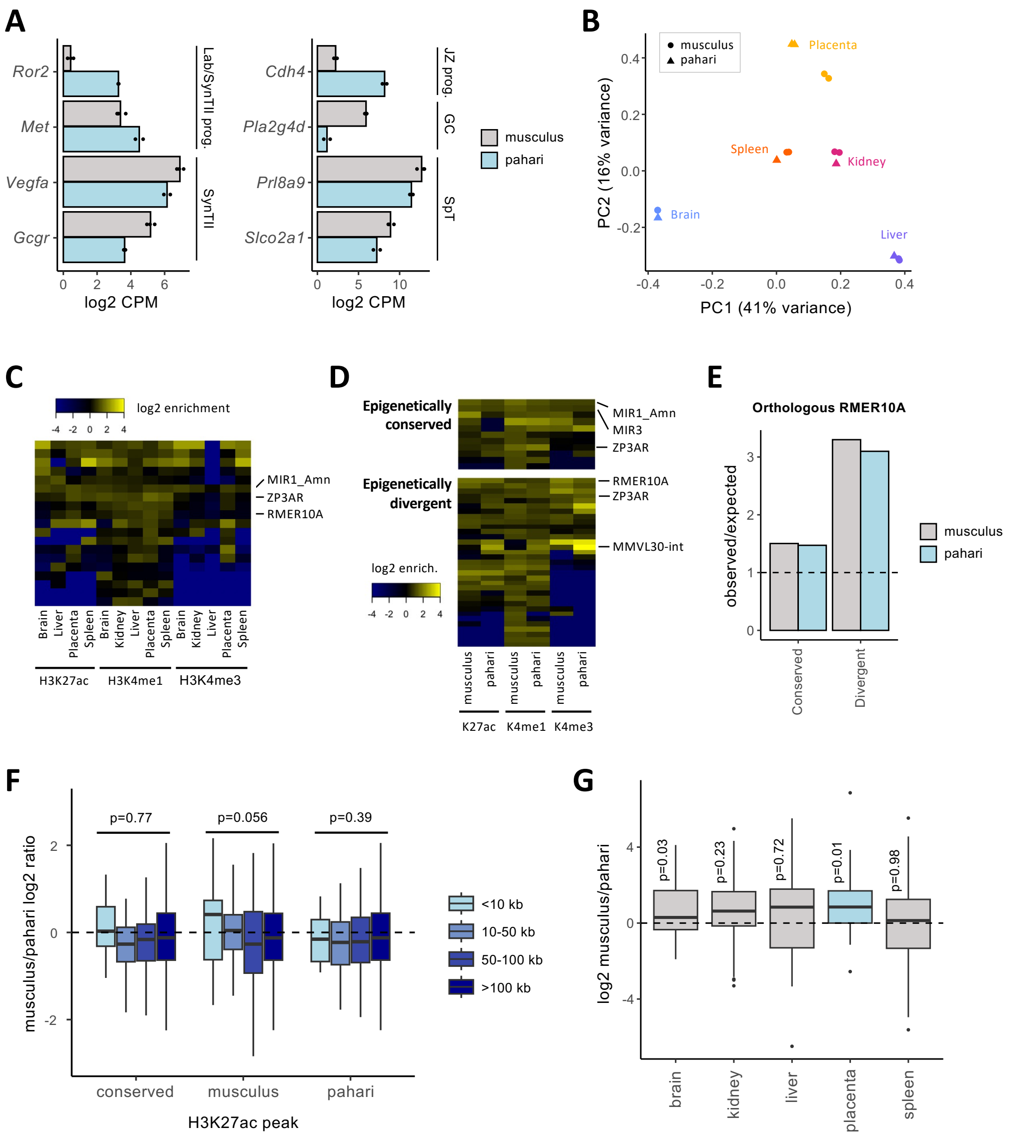


**Figure S8 – Comparative analysis of *musculus* and *pahari* placentas.** A) Markers of trophoblast cell types that are differentially expressed between *musculus* and *pahari* placentas (Lab – labyrinth, SynT – syncytiotrophoblast, JZ – junctional zone, GC – glycogen cells, SpT – spongiotrophoblast, prog. – progenitors). B) Principal component analysis of RNA-seq data from different *musculus* and *pahari* tissues. C) TE subfamilies displaying enrichment (observed/expected) for elements overlapping H3K27ac, H3K4me1 or H3K4me3 peaks in at least one *pahari* dataset. D) Enrichment at orthologous TEs for histone modification peaks that are present in both species (epigenetically conserved) or only in one (epigenetically divergent). E) Enrichment at orthologous RMER10A elements for epigenetically conserved or divergent H3K27ac peaks. F) Relative expression (*musculus*/pahari) of genes containing TSSs within the indicated distances from RMER10A elements with H3K27ac in the placenta of *musculus*, *pahari*, or both (conserved). G) Relative expression (*musculus*/pahari) of genes within 10 kb of an H3K4me3-marked IAP in *musculus* placenta.

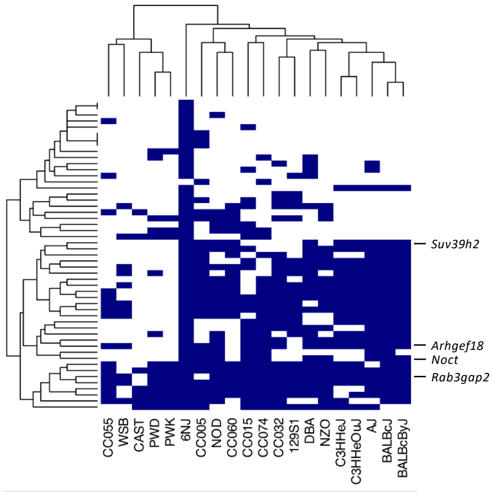


**Figure S9 – IAP insertional polymorphism across mouse strains.** Panel indicating the presence (blue) or absence (white) of H3K27ac-marked IAPs in different mouse strains. The genes annotated on the right contain isoforms incorporating that respective IAP.
